# Broad scale environmental conditions, not urbanisation, explain geographic variation in great tit feather microstructure

**DOI:** 10.64898/2026.09.21.753101

**Authors:** David López-Idiáquez, Hannah Watson, Javier Pérez-Tris, Caroline Isaksson, Pablo Salmón

## Abstract

Urbanisation imposes novel environmental conditions that can shape phenotypic traits including plumage, yet, its effects on characteristics beyond colouration remain poorly understood. Feather microstructure plays an important role in thermoregulation, protection from solar radiation and waterproofing, and its expression is sensitive to environmental conditions. Here, we quantified feather barbule density, a key component of feather microstructure, in great tits (*Parus major*) from five urban-rural paired populations across Europe to assess its association with urbanisation, temperature and elevation. We found no significant effects of the urban environment or temperature. However, feather microstructure significantly varied between the studied areas and was positively associated with elevation and ultraviolet B (UV-B) radiation, suggesting a potential adaptive response to variation in solar radiation.

## Introduction

Research on urbanisation as a phenotypic driver has revealed changes in traits such as phenology (Capilla-Lasheras et al. 2022, Cuchot et al. 2026), ornamentation (Halfwerk et al. 2018, Janas et al. 2024) and physiology (Salmón et al. 2018, Isaksson and Bonier 2020). Although relatively new ecosystems, urban environments impose novel and strong selective pressures on resident wildlife through factors such as pollution, reduced food quality/availability and temperature (Slabbekoorn and Peet 2003, Isaksson 2010, Szulkin et al. 2020). Importantly, these pressures are usually city specific, and populations can exhibit divergent responses at the phenotypic and genetic levels depending on the urban area (Alberti et al. 2020, Capilla-Lasheras et al. 2022). For instance, studies comparing urban and rural great tits (*Parus major*) across Europe have revealed polygenic city-specific responses to urban life (Salmón et al. 2021, Caizergues et al. 2022). Therefore, understanding population responses to urbanisation requires replicated studies across multiple urban populations to distinguish general patterns from city-specific effects. This is a particularly pressing given the predicted global expansion of urban areas (Humbal et al. 2023), which will amplify the impact of these novel selective pressures across populations.

In recent years, research on phenotypic responses to urbanisation has increasingly focused on avian plumage (e.g. (Amaya-Mejia et al. 2025, Janas et al. 2026, Bekka et al. 2026). Feathers are keratinous skin derivatives involved in a plethora of functions that, due to gradual abrasion and wear, must be periodically replaced during moult (Jenni and Winkler 2020). This feather replacement process is a costly activity that entails synthesising up to one third of the protein mass of a bird’s body (Murphy et al. 1988, Jenni and Winkler 2020) and, in many cases, adding costly pigments such as carotenoids (Thomas et al. 2014). These high production costs imply that feather quality can be, to a certain extent, environmentally sensitive and therefore, constrained in urban environments. This is, in fact, the case of plumage colouration that has recently been under close scrutiny, with several studies documenting consistent differences between urban and rural populations across space and time (Salmón et al. 2023, Janas et al. 2024, Yu et al. 2024, Sandmeyer et al. 2025, Bekka et al. 2026). However, while urban effects on colouration have received most of the attention, the effects on other plumage characteristics remain poorly understood. Among these, feather microstructure, the fine-scale organisation of feather barbs and barbules, is of particular relevance, as it plays a role in processes such as thermoregulation, protection from solar radiation and waterproofing (Terrill and Shultz 2023).

Experimental and correlative evidence has shown that feather microstructure is plastic and can respond to environmental conditions, e.g. food availability or temperature (Broggi *et al*. 2011, Gamero *et al*. 2015, Koskenpato *et al*. 2016). For instance, great tits inhabiting northern latitudes develop denser feathers than their southern counterparts (Broggi et al. 2011). Yet, whether the specific conditions characterising urban environments similarly shape feather microstructure has received comparatively less attention. To the best of our knowledge, only one study has addressed this question. Sándor et al. (2022), found that urban juvenile, but not adult, great tits developed feathers with lower barb density in two urban-rural population pairs in Hungary, likely reflecting reduced food availability and energy constraints during moult (Seress et al. 2020). Given the evidence of city-specific responses to urbanisation (Salmón et al. 2021, Caizergues et al. 2022) and geographic variation in feather microstructure (Gamero et al. 2015), replicated comparisons across population is required before the patterns reported by Sándor et al. (2022) can be generalised. The overarching aim of this study is to quantify the differences in feather microstructure between urban and rural great tits from five population pairs across Europe. Specifically, we aim to test whether urban great tits differ in their feather microstructure when compared with their rural counterparts. Furthermore, because of the multipopulational framework of our study, we also aim to analyse among-population differences in the feather microstructure and their environmental drivers. We predict that urban great tits will have less developed microstructure relative to rural birds, and that these structural differences will significantly vary across the five studied geographical areas.

## Methods

### Data sampling and feather measurement

Between 2014 and 2015, we captured 208 juvenile (1yr) and adult (2+ yr) great tits in five paired urban-rural habitats across Europe. At the time of capture and following a standardised protocol, we sampled a total of eight breast feathers per individual and stored them in paper envelopes until time of measurement.

We estimated feather microstructure (i.e. barbule density) in a subsample of three randomly selected feathers per individual using a semi-automated method applied to three barbs per feather. With that aim, we took a picture of each feather using a Dino-Lite microscope (AM7915MZT) at a fixed magnification. We then used ImageJ to convert the images to greyscale and extracted the intensity profile across three sections of approximately 1 mm, each drawn perpendicular to a different barb. Finally, we used a custom R function (Kiss 2024) to count the number of local minima in each profile, with each minimum corresponding to a single barbule (for further information, see SM-1). We estimated the technical repeatability of this method by manually counting the barbules in a subset of 84 barbs. Repeatability was high (0.89 [0.85, 0.93]), supporting the robustness of the semiautomated approach. Barbule density per barb was computed by dividing the number of barbules counted by the length of the measured section in millimetres. Then, we averaged the values of the three measured barbs in each feather, and this variable was used in the subsequent analyses.

To analyse the effects of temperature on feather microstructure, we obtained the temperature data at each location from the E-OBS Gridded Dataset v.31.0 with a resolution of 0.25 degrees (Cornes et al. 2018). Specifically, we computed mean temperature between 1-May and 31-August (mean temperature pre-moult) to capture the temperature experienced by the birds before and during moulting (Jenni and Winkler 2020). For Lisbon and Malmö, we used the temperature of the year of capture, as the birds were sampled after moult. For Madrid, Milan and Gothenburg, we used temperature data from the year before sampling, as those birds were captured before moulting. For UV-B radiation we used mean values between 2004 and 2013 in the same months as temperature, obtained from the glUV dataset (Beckmann et al. 2014), as a proxy of the UV radiation levels prevailing at each location.

### Statistical analysis

First, we examined differences in feather microstructure between urban and rural environments. With that aim, we fitted a linear mixed effects model (LMM) that included barbule density of each feather as the dependent variable. Habitat (urban vs. rural), sex and age (1 vs. 2+ years), and the interaction between habitat and age were included as fixed effects. Sampling area, population and individual identity were included as random effects to control for the non-independence of the samples taken in the same sampling area (n=10), the same population (n=5) and the same individual (n=208).

Second, we tested whether the effect of urbanisation on barbule density varied across populations. With that aim, we fitted an LMM including barbule density as the dependent variable. As explanatory terms, we included habitat, population, sex and age. As random effects, we included sampling area and individual identity.

Third we analysed the association between barbule density, elevation, temperature and UV-B radiation by fitting two separate LMMs. We first analysed the association with elevation in a model that included barbule density as the dependent variable and elevation, sex, and age as fixed effects. Then, to provide a mechanistic explanation for the potential links between barbule density and elevation, we fitted a second LMM that included barbule density as the dependent variable and temperature, UV-B radiation, sex, and age as fixed effects. Both models included sampling area, population and individual identity as random effects

To analyse temperature and elevation effects on barbule density, we fitted two separate LMMs including barbule density as the dependent variable and mean pre-moult temperature and elevation as the explanatory terms. In both models sex and age were included as covariates and sampling area, population and individual identity as random effects.

All models were fitted in R v. 4.5.2 (R Core Team 2023) using the packages *lme4* v. 1.1-37 (Bates et al. 2015) and *lmerTest* v. 3.1-3 (Kuznetsova et al. 2017). Model residuals were visually inspected and showed no marked deviations from normality. The significance of the model was estimated using *F-tests* based on the Satterthwaite approximation in *lmerTest*. Non-significant interactions (p>0.05) were removed from the models. Pairwise comparisons between factor levels were conducted using Tukey HSD correction with the package *emmeans* v. 2.0.0 (Lenth 2023).

## Results

Barbule density did not differ between urban and rural great tits, neither in adults nor in juveniles (Table 1, Fig. 1A). This lack of differences in barbule density between habitats was consistent across the studied areas (Table 2, Fig. 1B). However, the results from that model revealed significant between population differences: great tits from Madrid showed plumages with denser feathers than great tits from Malmo, Milan and Lisbon but not from Gothenburg (Fig. 2B).

**Table 1.**
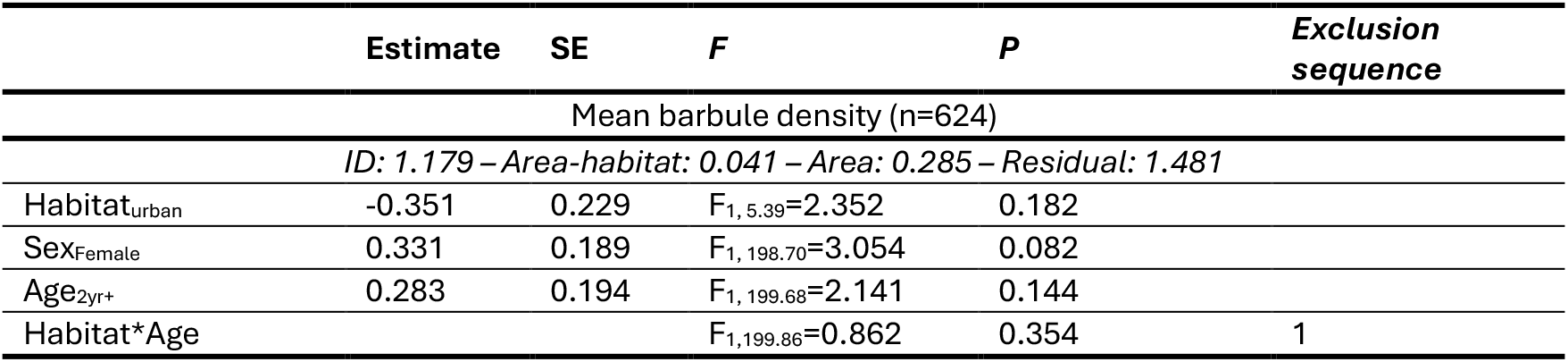
Results from the linear mixed-effects models analysing the difference in mean barbule density between urban and rural habitats. Estimates from the isolated terms and variance explained by random effects were obtained from the model not including the interaction.

**Table 2.** Results from the linear mixed-effects models testing for differential urban effects across geographic areas. Estimates from the isolated terms were obtained from the model not including the interaction. Bold denotes statistical significance.

|  | Estimate | SE | F | P | Exclusion sequence |
| --- | --- | --- | --- | --- | --- |
| Mean barbule density (n=624) |  |  |  |  |  |
| *ID: 1.198 – Residual: 1.482* |  |  |  |  |  |
| Habitat<sub>urban</sub> | -0.281 | 0.183 | F<sub>1, 200</sub>=2.394 | 0.126 |  |
| Sex<sub>Female</sub> | 0.330 | 0.190 | F<sub>1, 200</sub>=3.003 | 0.084 |  |
| Age<sub>2yr+</sub> | 0.265 | 0.193 | F<sub>1, 200</sub>=1.883 | 0.171 |  |
| **Area** |  |  | **F<sub>4, 200</sub>=9.333** | **<0.001** |  |
| Habitat*Area |  |  | F<sub>4, 196</sub>=1.256 | 0.288 | 1 |

**Figure 1.**
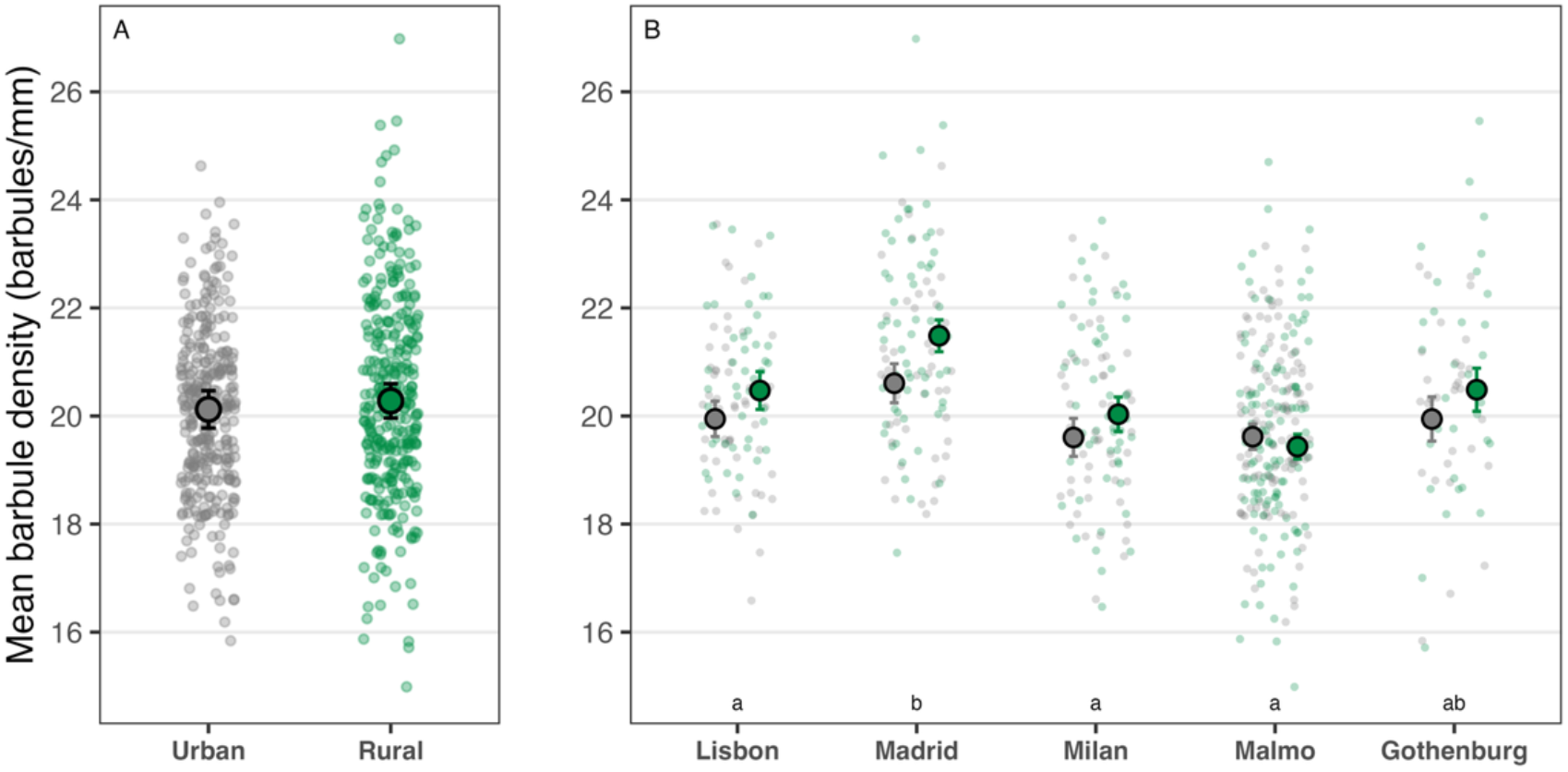
Effects of urbanisation on mean barbule density. A, represents the overall differences between the urban and rural environments. B, represent the differences between the urban and rural environments in each geographic area. Dots and whiskers represent the mean predicted barbule density ± standard error. Grey and green dots in the background represent the raw observations from the urban and rural habitats respectively. In B, letters over each population denote statistical differences in barbule density between areas. Areas with no letters in common are significantly different (p<0.05).

**Figure 2.**
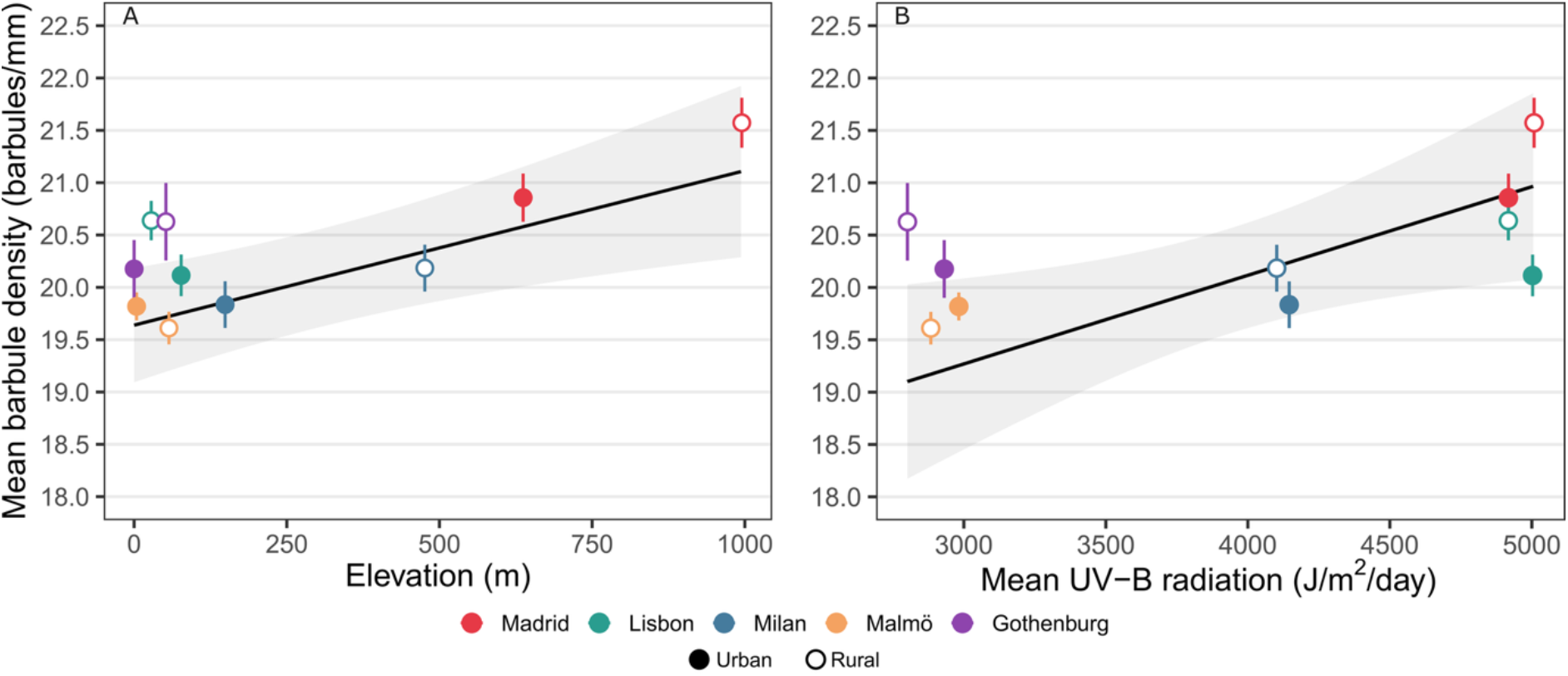
A) Association between elevation (meters above sea level) and mean barbule density. B) Association between mean UV-B radiation and mean barbule density. The black lines and ribbons represent the trend predicted from the model, and the dots and whiskers represent the mean barbule density in each sampling area ± standard error.

We found a significant positive association between mean barbule density and elevation, with birds inhabiting higher elevation populations bearing more dense feathers (0.001±0.0004 barbules m^−1^, F_1,6.593_=9.122, p=0.020; Fig. 2A, see SM-2 for full model results). Evaluating the potential environmental drivers of this association, temperature was not associated with barbule density (F_1,4.296_=2.289, p=0.159), whereas birds inhabiting in areas with more intense UV-B radiation had denser feathers (0.0008±0.0003, F_1, 6.925_=5.828, p=0.046, Fig. 2B, see SM-2 for full model results).

We found a significant positive association between mean barbule density and elevation, with birds inhabiting higher elevation populations bearing more dense feathers (0.001±0.0004 barbules m^−1^, F_1,6.593_=9.122, p=0.020; Fig. 2A, see SM-2 for full model results). Testing the potential environmental drivers of this association, temperature was not associated with barbule density (F_1,4.296_=2.289, p=0.159), whereas birds inhabiting in areas with more intense UV-B radiation had denser feathers (0.0008±0.0003, F_1, 6.925_=5.828, p=0.046, Fig. 2B, see SM-2 for full model results).

## Discussion

Contrary to our first prediction, we found no evidence for differences in feather microstructure between urban and rural great tits in five replicated population pairs across Europe. Our results did, however, support our second prediction, as we found significant variation among the five studied areas. Our analyses show that the between population variability was, to a certain extent, explained by elevation with great tit populations at higher elevations having feathers with denser microstructures. Linking feather micro-structure to two altitude-associated environmental factors, UV-B radiation and temperature, indicates that the between-population differences can, to a certain extent, be explained by UV-B radiation, a result that provides a mechanistic explanation for the links with elevation. Temperature, on the other hand, was not significantly associated with feather microstructure. Overall, this suggests that urban characteristics do not strongly influence feather microstructure, which is instead more driven by other macroecological and climatic factors.

The lack of a widespread urban effect on adult feather microstructure reported here contrasts with the only other published study tackling a similar question. Sándor *et al*. (2022) reported reduced barbule and barb density in contour and wing feathers of urban juvenile (i.e. 1-year-old), but not adult, great tits. Explaining the differences between that study and our results is not an easy task, but it may be rooted in the different geographical coverage of the two studies. While Sándor *et al*. (2022) focus on two closely located urban-rural pairs (i.e. in the same region in Hungary), we analysed five population pairs spanning from northern to southern Europe. Thus, it is likely that, in the two sets of populations, the potential pressures imposed by urbanisation on feather microstructure differ. The absence of a significant difference between urban and rural great tit feather microstructure described here also contrasts with previous research reporting urban effects on other plumage characteristics (Sándor *et al*. 2021, Salmón *et al*. 2023, Sandmeyer *et al*. 2025, but see Amaya-Mejia *et al*. 2025). For instance, Salmón et. al (2023) reported paler colouration in urban great tits when compared with their rural counterparts. As they used the same multi-population framework as in the present study, this suggests that our analytical set up is robust to the detection of urban-driven phenotypic shifts, thus supporting the observed lack of an urban effect on feather microstructure in the present study. Previous research has also reported urban effects on other feather characteristics with, for instance, urban birds developing a reduced number of feathers than their rural counterparts (Sándor et al. 2021). Interestingly, most of these studies report significant urban effects in juveniles but not in adults. Taken together, these results led us to suggest that feather microstructure, unlike other plumage characteristics, is not consistently impacted by the environmental stressors associated with urban environments and may be modulated by other ecological factors.

The existence of between-population differences in feather microstructure are in line with previous research in great tits (Broggi et al. 2011, Gamero et al. 2015). Because feather microstructure is highly plastic, these broad population differences probably reflect geographical variation in environmental conditions (Gienapp and Merilä 2010, Broggi et al. 2011). Given that thermoregulation is one of the main plumage functions (Terrill and Shultz 2023), ambient tem-perature has often been identified as a key factor explaining variation in feather structure within (Koskenpato et al. 2016, Nord et al. 2023, Amaya-Mejia et al. 2025, González-Medina et al. 2026) and among species (Pap et al. 2017). For instance, in tawny owls (*Strix aluco*), differences in feather microstructure between brown and grey morphs is one of the proposed explanations for the reduced overwinter survival of brown-morph individuals during harsh winters (Koskenpato et al. 2016). Interestingly, our results for great tits do not support this idea as we did not find a significant association between temperature and feather microstructure. This absence of sensitivity to variation in temperature aligns with previous evidence in other great tit populations (Gamero et al. 2015, Sándor et al. 2022). For instance, great tits from Lund (southern Sweden) exhibit higher barbule densities than great tits from Barcelona (Spain) and Oulu (northern Finland) despite the latter experiencing less extreme temperatures (Gamero et al. 2015). We found a positive association between feather microstructure and elevation. Although few studies have examined this specific relationship, the existing evidence links elevation to feather structure both within and between species (Cheek et al. 2018, Barve et al. 2021). For instance, torrent ducks (*Merganetta armata*) at lower altitudes have shorter down with fewer barbs compared with conspecifics living at higher altitudes (Cheek et al. 2018). In most cases, these associations have been explained by the superior insulation benefits of feathers with more developed microstructure. This explanation, however, does not fit our results, given that we explicitly tested the effects of temperature, rather than using elevation as a surrogate, without finding any significant association. We propose instead that the observed altitudinal trend might be driven by solar radiation as is supported by our results linking feather structure and UV-B radiation. Feather microstructure is tightly linked with the ability of the feathers to reflect both avian-visible (300-700 nm) and the near-infrared (NIR; 300-2600 nm) wavelengths of sunlight (Stuart-Fox et al. 2018). In addition to its thermal effects, solar radiation can also have other pervasive effects on birds, such as cellular damage, immunosuppression or feather degradation (Blount and Pike 2012, Galván et al. 2026). For instance, experimental studies have shown that sunlight ultraviolet (UV) radiation can impair pro-inflammatory immune responses in zebra finches (*Taeniopygia guttata*) (Blount and Pike 2012) and that solar photobleaching leads to feather degradation in Spanish imperial eagle (*Aquila adalberti*) fledglings, potentially limiting flight performance (Galván et al. 2026). Thus, we suggest that the more developed feather microstructure at higher elevations represents an adaptive response to solar radiation, given the expected increase in solar radiation intensity with elevation, arising from reduced atmospheric attenuation at higher altitudes. Nonetheless, future studies sampling an altitudinal gradient within a single site, rather than across geographically distinct populations, are required to corroborate this finding.

## Conclusion

Using a replicated sampling design of five urban-rural great tit population pairs across Europe, we found no significant effects of urbanisation on feather microstructure. Our results, however, showed between-population differences in this feather trait, which were largely explained by elevation, with populations at higher altitudes having denser micro-structures. We suggest that this structural shift may represent an adaptive response to the increased solar radiation at higher altitudes. Between-population variation in temperature was not associated with variation in feather micro-structure contrasting with studies in other species and suggesting that in great tits feather microstructure may not play a major role in thermoregulation.

## Acknowledgements

We thank everyone who assisted during the feather sampling, including Andreas Nord, ringers from La Herrería, Jose Ignacio Aguirre, Michelangelo Morganti and Lago di Pusiano ringers, Vítor Encarnação and the Pardal family. We also thanks Anett Kiss for her help with the semi-automated counting method. Finally, we acknowledge the E-OBS dataset from the EU-FP6 project UERRA (https://www.uerra.eu) and the Copernicus Climate Change Service and the data providers in the ECA&D project (https://www.ecad.eu).

## Data availability statement

All data and code required to replicate the analyses presented in this manuscript are available on GitHub (https://github.com/idiaquez2/Urban-Effects-on-FeatherStructure.git)

