## Supplementary Material for "Broad scale environmental conditions, not urbanisation, explain geographic variation in great tit feather microstructure"

**SM-1 Further details on the process followed to count the number of barbules.**

Feather pictures were taken using a Dino-Lite microscope (AM7915MZT). The distance between the feather and the microscope, and the microscope's magnification, were kept constant in every picture. Thus, in all pictures 1 mm corresponded to the same number of pixels (329 pixels; SM-1 Fig. 1A). These pictures were converted to greyscale using ImageJ, and local contrast was enhanced to maximise the contrast between the feather structures and the background (SM-1 Fig. 1B). Then, using ImageJ‘s straight-line tool, we drew three straight lines perpendicular to three different barbs. Finally, we exported the intensity profile across each line using the multi-plot tool in ImageJ (SM-1 Fig. 1C). These profiles were analysed using a custom R function (Kiss 2024) to count the number of local minima, which represent individual barbules crossing each line (i.e. the dark troughs in the intensity profile, with peaks representing the spaces between barbules). The custom function computed the number of barbules per 1 mm (i.e. barbule density), accounting for the fact that line length was slightly variable (mean ± SD: 1.01 ± 0.014 mm).

We validated this approach by manually counting barbules in a subset of 28 feathers (84 barbs). The mean absolute difference between manual and automated counts was ±0.583 barbules. We also assessed the agreement between the two approaches by computing repeatability using the package rptR (v. 0.9.23, Stoffel *et al.* 2017), both at the barbule and feather level. In both cases repeatability was significant and high: 0.898 [0.85, 0.93] and 0.703 [0.53, 0.80], respectively. The small absolute differences and high repeatability together support the reliability of our automated approach.

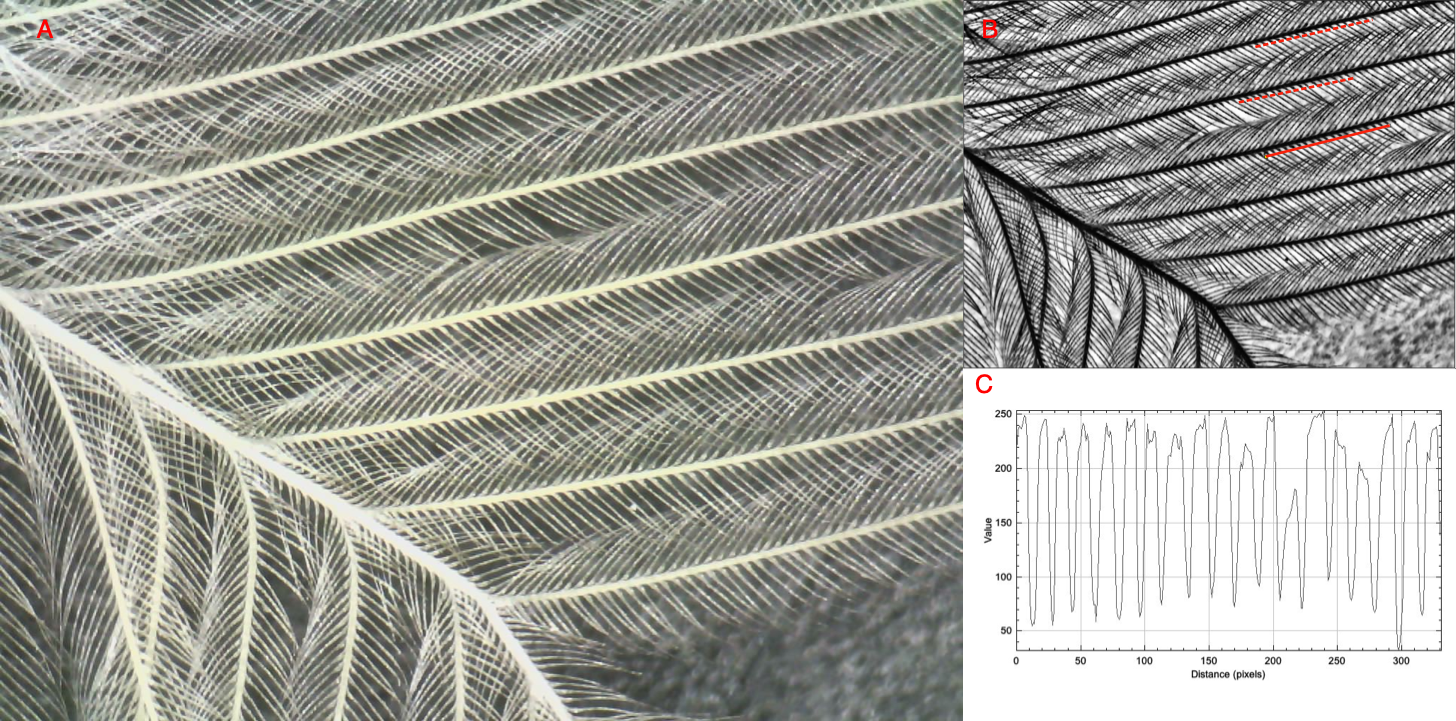

**Figure SM-1:** Schematic representation of the process to count the number of barbules in each feather. A represents the original picture of one of the measured feathers, and B represents the same picture but in black and white. In B, the red lines represent the sections where the number of barbules were counted. C represents the change in colour pattern across the solid line; here, local minima represent each barbule. Note that red lines are displayed for illustration purposes and are not the actual lines drawn in ImageJ.

**SM-2 Full model results**

**SM-2 Table 1:** Results from the linear mixed-effects models analysing the link between elevation and feather microstructure. Bold denotes statistical significance.

|  | **Estimate** | **SE** | ***F*** | ***P*** |
| --- | --- | --- | --- | --- |
| Mean barbule density (n=624) | | | | |
| *ID: 1.183 – Area-habitat: <0.0001 – Area: 0.114 – Residual:1.482* | | | | |
| Sex_Female_ | 0.321 | 0.189 | F_1, 202.26_=2.898 | 0.090 |
| Age_2yr+_ | 0.289 | 0.192 | F_1, 203.93_=2.281 | 0.132 |
| **Elevation** | **0.001** | **0.0004** | **F_1, 6.593_=9.122** | **0.020** |

**SM-2 Table 2:** Results from the linear mixed-effects models analysing the link between mean temperature in May-August, ultraviolet (UV)-B radiation and feather microstructure. Bold denotes statistical significance.

|  | **Estimate** | **SE** | ***F*** | ***P*** |
| --- | --- | --- | --- | --- |
| Mean barbule density (n=624) | | | | |
| *ID: – Area-habitat: – Area: – Residual:* | | | | |
| Sex_Female_ | 0.331 | 0.190 | F_1, 196.70_=3.042 | 0.082 |
| Age_2yr+_ | 0.283 | 0.195 | F_1, 199.90_=2.101 | 0.148 |
| Mean temp. | -0.199 | 0.117 | F_1, 4.296_=2.289 | 0.159 |
| **UV-B** | **0.0008** | **0.0003** | **F_1, 6.925_=5.828** | **0.046** |
